# Urgency disrupts cognitive control of human action

**DOI:** 10.1101/2021.09.08.459465

**Authors:** Christian H. Poth

**Author notes:** Correspondence should be addressed to*: Christian Poth, Department of Psychology, Bielefeld University, P.O. box 10 01 31, D-33501 Bielefeld, Germany.

## Abstract

Intelligent behavior requires cognitive control, the ability to act directed by goals despite competing action tendencies triggered by stimuli in the environment. For eye movements, it has recently been discovered that cognitive control is briefly disrupted in urgent situations (Salinas et al., 2019). In a time-window before an urgent response, participants could not help but look at a suddenly appearing visual stimulus, even though their goal was to look away from it. Urgency seemed to provoke a new visual-oculomotor phenomenon: A period in which saccadic eye movements are dominated by external stimuli, and uncontrollable by current goals. This period was assumed to arise from brain mechanisms controlling eye movements and spatial attention, such as those of the frontal eye field. Here, we show that the phenomenon is more general than previously thought. We found that urgency disrupted cognitive control also in well-investigated manual tasks, so that responses were dominated by goal-conflicting stimulus features. This dominance of behavior followed established trial-to-trial signatures of cognitive control that replicate across a variety of tasks. Thus together, these findings reveal that urgency temporarily impairs cognitive control in general, not only at brain mechanisms controlling eye movements.

## Introduction

Intelligent behavior requires that information about the environment is not only processed, but that it is reconciled with ongoing actions and momentary behavioral goals. This requires *cognitive control,* the ability to behave goal-directed even if this means to overcome compelling tendencies for other behaviors (Cohen, 2017; Gratton et al., 2018). Cognitive control generally relies on brain mechanisms of the frontal cortex (Miller, 2000; Nee and D’Esposito, 2016; Ridderinkhof, 2004; Tang et al., 2016), which is evident from the devastating impairments of cognitive control in neurological disorders with frontal damages (Alexander et al., 2007; Badre et al., 2009; Casey et al., 2002). These brain mechanisms are thought to interface with attention mechanisms at multiple levels of processing (e.g., Egner, 2008), thereby enforcing that goal-relevant information is prioritized, whereas conflicting information is rejected (Cohen, 2017; Gratton et al., 2018).

Salinas et al. (2019) found that the need for urgent responding opened up a time-window (~40 ms) in which the cognitive control of saccadic eye movements was severely reduced. Participants had to look away from a suddenly appearing visual stimulus (an antisaccade), which requires to suppress the natural tendency to look toward it (Munoz and Everling, 2004). During the time-window, Salinas et al. observed that the eye movements were strongly drawn to the location of the visual stimulus, even though this conflicted with current task-goals. The authors interpreted this finding as evidence for an “attentional vortex”, a time-window in which the endogenous (goal-directed) control of spatial attention was overpowered by the exogenous attraction of spatial attention to the stimulus location. However, this interpretation in terms of attention control was hotly debated during eLife’s open peer-review (https://elifesciences.org/articles/46359#SA1). Given the tight links of vision and eye movements, it was suggested that the findings reflected a specific visually-driven impairment of eye movement control (oculomotor capture; e.g., Irwin et al., 2000), and that “the link to attention is only inferred” (review round 2, point 1). Compatible with the suggestion, Salinas et al. modeled potential mechanisms underlying their findings based on neuronal activity of brain regions specialized in eye movement control (Costello et al., 2013; Seideman et al., 2018; Shankar et al., 2011; Stanford et al., 2010). Thus, in line with the suggestion, this modeling constrains interpretations to the domain of eye movements and closely related brain mechanisms of spatial attention (Moore and Armstrong, 2003).

The critical debate urges us to ask how general Salinas et al.’s discoveries are. Specifically, are the findings really limited to eye movement control and its linked attention mechanisms, or do they signify more wide-ranging impairments of cognitive mechanisms? We show that the latter is the case. Going even beyond Salinas et al.’s “attentional vortex”, we found that urgency leads to a temporary loss of cognitive control in general. In two well-established manual tasks of cognitive control, behavioral responses were dominated by task-irrelevant information. As a result, participants could not help but execute responses that were in clear conflict with their current goals. Experiment 1 revealed such a loss of cognitive control for conflicts of spatial stimulus location, and Experiment 2 demonstrated the same pattern even for non-spatial cognitive conflicts. In both experiments, the performance drops followed trial-to-trial sequence effects that signify top-down and/or bottom-up modulations of cognitive control (Egner, 2017, 2007). As such, these findings offer mounting evidence that urgency temporarily disrupts cognitive control of human action at levels extending beyond vision, spatial attention, and eye movement control (such as prefrontal brain mechanisms; Egner, 2007; Egner and Hirsch, 2005).

## Results

### Urgency temporarily disrupts cognitive control of spatial conflicts

In Experiment 1, cognitive control was assessed using a spatial Stroop task (Clark and Brownell, 1975; Funes et al., 2010; Lu and Proctor, 1995; Schneider, 2020). In this task, participants had to respond by pressing one of two buttons to indicate whether a target stimulus, an arrow (see Figure 1), was pointing left or right. The target arrow could appear either to the left or right of screen center. In the congruent condition, the target arrow appeared on the side it was pointing to. In the incongruent condition, the target arrow appeared on the side opposite to where it was pointing to. Such a condition has been shown to lead to slower reaction times, indicating that the spatial location and the pointing direction of the target arrow produced a cognitive conflict for action (Lu and Proctor, 1995; Schneider, 2020, Exp. 1). Cognitive control can be quantified by assessing how well this cognitive conflict can be resolved. This is done by measuring how performance suffers in the incongruent condition compared with the congruent condition. If cognitive control was perfect, performance should be equal in the conditions, and for worse cognitive control, performance should drop stronger in the incongruent compared with the congruent condition. Harnessing these classic congruency effects, we traced the evolution of cognitive control over time using a manipulation of urgency. The trial sequence is shown in Figure 1. In the beginning of a trial, participants fixated a small square in the center of the screen. The disappearance of this fixation stimulus served as the go-signal, prompting participants to respond after one second. Urgency was manipulated by introducing a gap of different durations after the go-signal and before the target stimulus that indicated the response to be made (Salinas et al., 2019; Stanford et al., 2010). When the duration of the gap is short, urgency should be low. The target appears just after the go-signal, allowing for processing the target in order to prepare the motor responses. In contrast, when the gap duration is long, urgency should high. During the gap after the go-signal, the time left for responding is elapsing, but the target specifying the response is still missing. Thus, time-pressure for responding is building up, so that participants must prepare their motor responses before it becomes clear which response has to be chosen. This manipulation induced urgency as an internal state of participants’ cognitive system, which was evident from increases in physiological arousal with increasing gap duration (*supplemental materials, Figure S2*). Following Salinas et al. (2019), we computed the raw processing time (RPT) for the target as the time the target was visible within reaction time (i.e. the reaction time from the go-signal minus the gap duration). For both conditions, we assessed the temporal evolution of performance by computing the tachometric function, which is the psychometric function relating performance to RPT (*Figure 2a*). This revealed a qualitative difference between the congruent and incongruent condition (and closely replicated a pilot experiment, see the *supplemental materials, Figure S1*). In the congruent condition, performance rose monotonously from chance level up to an asymptote at near-perfect performance. In contrast, the tachometric function in the incongruent condition followed a qualitatively different course: While it also started at chance and evolved toward near-perfect performance, it dropped below chance level in a demarcated time-window around an RPT of about 200 ms. The maximum drop below chance was lower in the incongruent condition than the congruent condition (*p* < .001, permutation test, see methods; *Figure 2b*). This reveals that there is a demarcated time-window of RPTs, in which cognitive control is severely impaired, so that against task-instructions, responses are dominated by the spatial location of the target arrow rather than its pointing direction.

**Figure 1.**
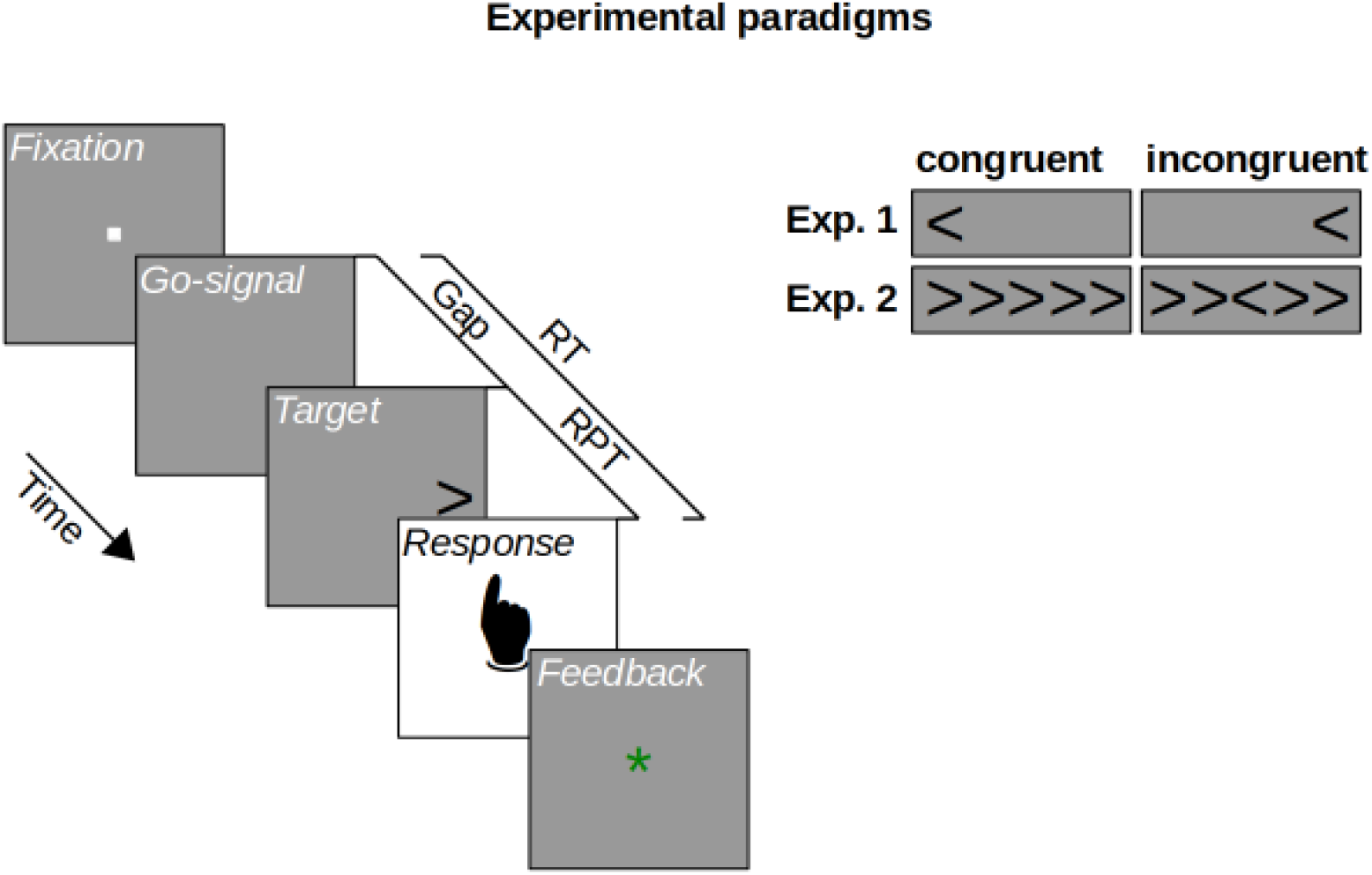
Experimental paradigms. On each trial, participants fixated a fixation stimulus, whose disappearance served as a go-signal, prompting a response within a 1 s deadline. In both experiments, the target stimulus followed the go- signa after a variable gap duration. Shorter gaps imply lower urgency for responding, leaving enough time until the deadline. Longer gaps imply higher urgency for responding, leaving little or no time until the deadline. Participants responded by button press, receiving a feedback about whether they responded in time. Experiment 1 used a spatial Stroop task, in which participants indicated the direction of an arrow matching or mismatching (congruent vs. incongruent) its horizontal location. Experiment 2 used a flanker task, in which participants indicated the direction of a central arrow flanked by matching or mismatching (congruent vs. incongruent) distractor stimuli. Raw processing time (RPT) is the reaction time (RT) minus gap duration.

**Figure 2.**
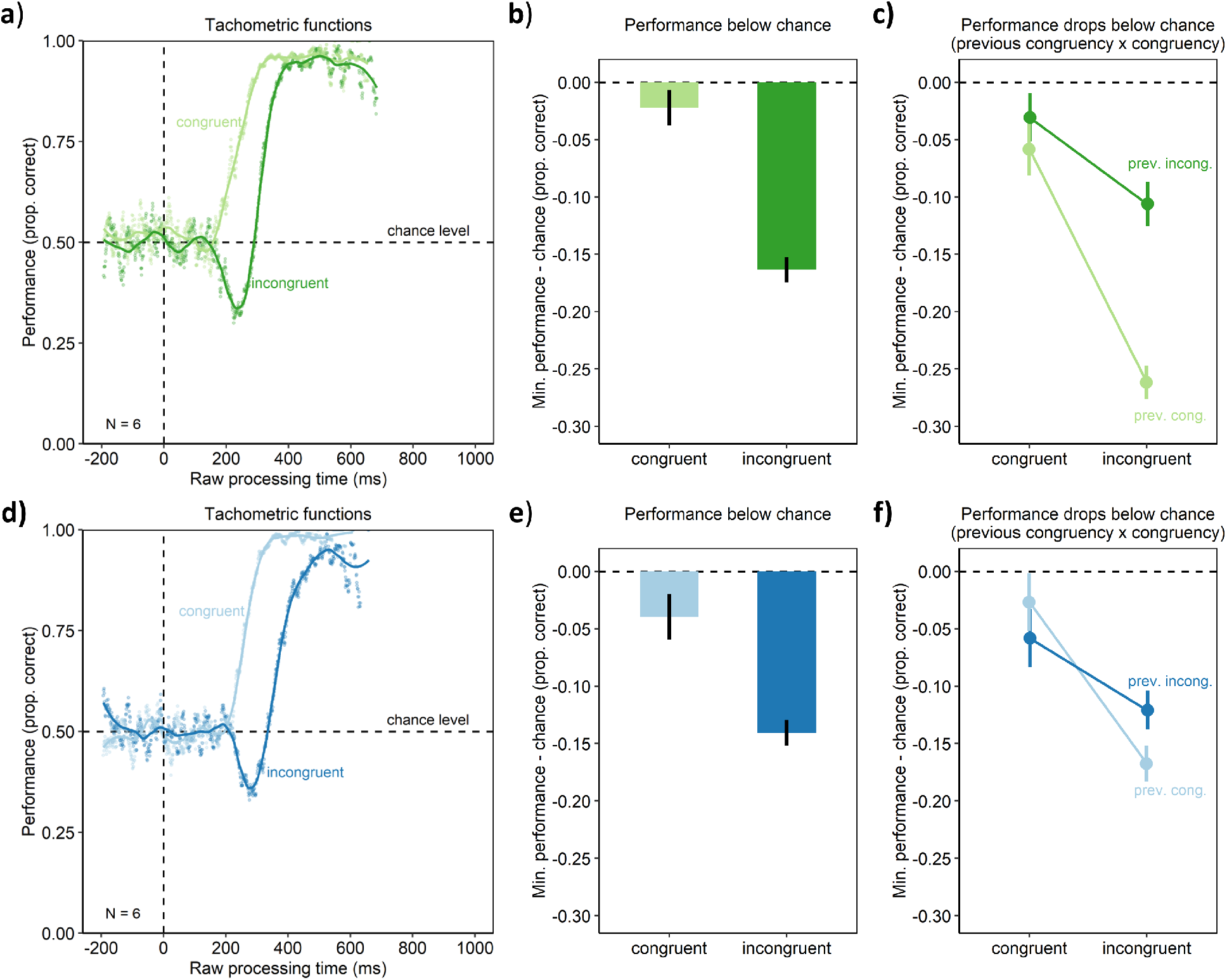
Behavioral results of Experiment 1 (upper panel) and Experiment 2 (lower panel). a/d) Tachometric functions depicting performance as a function of RPT for the congruent and incongruent conditions of the aggregate participant. b/e) In contrast to the congruent condition, the incongruent condition led to a drop of performance below chance. c/f) The drop of performance below chance was larger, when the previous trial was congruent (prev. cong.) rather than incongruent (prev. incong.). Error-bars are standard errors from a bootstrap.

One hallmark of cognitive control is that it is dynamic: Humans can adapt or learn to compensate cognitive conflicts (Botvinick et al., 2001; Egner, 2007; Hommel et al., 2004), so that performance decrements are less pronounced following trials with cognitive conflict (incongruent trials) than trials without such conflict. We discovered that this was also the case for our urgency-based effects: Performance dropped more strongly below chance level in the incongruent condition when the preceding trial was congruent rather than incongruent (*p* < .001, permutation test, *Figure 2c*). Thus, the drops of performance below chance exhibit the same trial-to-trial modulations of established congruency effects. Thus, this pattern of results offers converging evidence that the performance drops reflect a temporary collapse of cognitive control.

Taken together, the results of spatial Stroop task of Experiment 1 reveal a demarcated time-window along the urgency-axis in which cognitive control collapses, so that responses cannot be based on the symbolic information of the target arrow (i.e. its pointing direction), but are involuntarily (against task instructions) based on the spatial location of the target. Salinas et al. observed similar effects for saccadic eye movements, in which urgency led to a time-window in which saccades were involuntarily drawn towards rather than against as per the instructions a suddenly appearing target. Thus, the present findings suggest that collapses in cognitive control are not limited to eye movements captured by sudden stimuli. Rather, cognitive control is also impaired in manual decision tasks. In Experiment 1, cognitive conflicts arose from the task-irrelevant spatial location of the target stimulus. Thus, impairments of cognitive control could be limited to such spatial cognitive conflicts. Therefore, we performed Experiment 2 to test whether our present findings translated to non-spatial types of cognitive conflict.

### Urgency temporarily disrupts cognitive control of non-spatial conflicts

In Experiment 2, cognitive control was assessed using a flanker task (Eriksen and Eriksen, 1974; Ridderinkhof et al., 2021). The paradigm and trial sequence were similar to the one of Experiment 1. However, here the target arrow always appeared at screen center. The target was flanked by four distractor arrows. In the congruent condition, the distractors pointed in the same direction as the target arrow, so that there was no cognitive conflict. In contrast, in the incongruent condition, the distractors pointed into the opposite direction as the target, thus providing conflicting information as to which response had to be executed. In contrast to Experiment 1, as the target always appeared at screen center, the conflict was not spatial in the sense that it was due to the target’s spatial location. Results revealed the same pattern as in Experiment 1. Performance monotonously rose from chance to near-perfect performance with increasing RPT in the congruent condition (*Figure 2d*). In contrast, within a demarcated time-window of RPTs, performance dropped below chance level in the incongruent condition (p < .001, permutation test). Thus, even though the type of cognitive conflict was non-spatial, urgency resulted in a temporary collapse of cognitive control during which responses were dominated by distractors, contrary to the task-instructions. As in Experiment 1, this collapse was subject to conflict adaptation, which is the trial-to-trial modulation typical for cognitive control. That is, collapses were stronger after trials with no conflict (congruent trials) compared to trials with conflict (incongruent trials). Taken together, the results of Experiment 2 suggest that urgency elicits a circumscribed time-window in which cognitive control is diminished also for non-spatial cognitive conflicts.

## Discussion

Urgency elicited a time-window in which the ability to resolve cognitive conflicts was lost. During this time-window, participants responses were dominated by stimulus information, forcing them to execute the stimulus-driven responses against their current task-goals. This pattern of results closely resembles the effects of abruptly appearing visual stimuli on saccadic eye movements found by Salinas et al. (2019). However, the present findings reveal that these urgency effects have more wide-ranging implications than previously thought. Rather than causing a limited visual-oculomotor phenomenon, urgency seems to impair cognitive control in general.

Cognitive control was temporarily lost in two manual tasks, showing that this phenomenon not restricted to saccadic eye movements. Compared with eye movements, this effect appeared for longer RPTs, in line with the generally longer reaction times for manual movements compared with eye movements. We found the phenomenon for cognitive conflicts in which the spatial location of stimuli conflicted with their task-relevant features (Experiment 1). This is in line with Salinas et al.’s interpretation that urgency caused an irresistibly strong capture of saccadic eye movements by the visual stimulus (oculomotor capture; Theeuwes, 2014) in their antisaccade task. Given the tight links of eye movement control and spatial attention (Deubel and Schneider, 1996; Moore and Armstrong, 2003; Rolfs et al., 2011), such a capture should not only attract saccades (overt attention), but also covert spatial attention (without saccadic eye movements) prioritizing the target’s location for perception (Carrasco, 2011; Theeuwes, 2014). For Experiment 1, such a prioritization of stimulus location could have provoked manual responses that were compatible in their spatial organization (Lu and Proctor, 1995). This would explain the findings of Experiment 1 by assuming mechanisms similar to those implicated in Salinas et al.’s eye movement paradigm. Critically, however, in Experiment 2 we found the loss of cognitive control also for cognitive conflicts that were non-spatial. Here, urgency elicited a time-window in which distractor stimuli flanking a target stimulus involuntarily dominated participants’ responses. Furthermore, in both experiments, the loss of cognitive control was stronger after trials with conflict than trials without conflict. Although the precise nature of top-down and bottom-up influences underlying such modulations is still debated (Egner, 2017), for our present view it offers corroborating evidence that the stimulus dominance of participants’ responses indeed reflects a loss of cognitive control. Thus, taken together, the present findings reveal that urgency disrupts cognitive control not only in saccadic eye movements and spatial attention, but in goal-directed behavior more generally.

How could urgency induce such a general temporary loss of cognitive control? In an urgent task, the motor plans for the response alternatives are active after the go-signal for responding, even if the stimulus indicating which response should be made has not yet appeared (Salinas et al., 2019; Stanford et al., 2010). Thus, the motor plans are both active and compete, and this competition must be resolved “on-the-fly” using incoming visual information. We found that the time-pressure to respond, and thus urgency, was generally associated with physiological arousal measures (pupil size, Mathôt, 2018; saccade peak velocity, Di Stasi et al., 2013). This invites the hypothesis that urgency increases neuronal arousal (e.g., by modulating nor-epinephrine levels, Aston-Jones and Cohen, 2005), and by this, it could generally speed up cognitive information processing (Bundesen et al., 2015). This would shorten the time over which the competition between motor plans extends, causing information with stronger neuronal representations to almost immediately decide the competition in its favor. No time would be left for cognitive control mechanisms to counteract the influence of this information. This would be in line with Salinas et al.’s (2019) physiologically inspired computational model, which explains the temporary loss of cognitive control as a period in which faster exogenous signals already affect motor plans while slower endogenous signals have not yet been processed up to this level.

It has long been established that cognitive conflicts slow down responses in a variety of cognitive tasks (Lu and Proctor, 1995; Ridderinkhof et al., 2021). However, one may wonder whether this actually makes a difference for behavior outside the laboratory, in real-life. In most situations, it may have little consequences if a cognitive conflicts delays a goal-directed response for a couple of tens (or even hundreds) of milliseconds. However, the present findings suggest that urgency carries a real danger here. Rather than just delaying goal-directed behavior, it enforces stimulus-driven behaviors that are adversarial to the current goals. Thus, the resulting behavioral errors may be a source of accidents in a number of safety-critical situations such as traffic situations.

## Conclusions

Urgency temporarily disrupted the ability to handle cognitive conflicts. Behavior was involuntarily dominated by stimulus information, even though this information was adversarial to current goals. This disruption of cognitive control seems general in the sense that it is 1) extends beyond previous reports of impaired eye movement control (Salinas et al., 2019), and 2) occurs also in manual tasks, and 3) occurs irrespective of whether cognitive conflicts arise from spatial stimulus locations or non-spatial stimulus information. Going beyond previous suspicions (Salinas et al., 2019), urgency seems to elicit a general “attentional vortex in cognitive control”, during which behavior is controlled by the external world rather than internal goals.

## Materials and methods

### Participants

Six participants (aged 21 – 23 years) performed Experiment 1, and 6 (aged 22 – 25 years) performed Experiment 2. These sample sizes were chosen in advance, based on Salinas et al. (2019) and based on the pilot experiment that was replicated by Experiment 1 (see, the supplemental materials). They all reported (corrected-to-) normal vision and gave written informed consent before participating. The experiment followed the ethical regulations of the German Psychological Society (DGPs) and was approved by Bielefeld University’s ethics committee.

### Experimental setup and tasks

Participants performed the experiments in a dimly lit room. They viewed the pre-heated (Poth and Horstmann, 2017) computer screen (G90fB, ViewSonic, Brea, CA, USA) from a distance of 71 cm, while their right eye was tracked (Eyelink 1000, tower-mounted eye tracker, SR Research, Ottawa, Ontario, CA). They responded using a standard computer mouse. The experiments were programmed using the Psychophysics Toolbox 3 (3.0.12; Brainard, 1997; Kleiner et al., 2007) and Eyelink toolbox (Cornelissen et al., 2002) for MATLAB (R2014b; The MathWorks, Natick, MA).

Figure 1 illustrates an experimental trial. Participants fixated a central fixation stimulus (a 0.2×0.2° square, 85 cd/m^2^) for 350, 400, or 500 ms, whose subsequent disappearance prompted them to respond within 1 s, even if the target stimulus for the response had not yet been shown. After a variable gap duration (0, 100, 200, …, 900, or 950 ms), the target stimulus appeared. Participants responded by pressing the left or right mouse button with their left or right index fingers, respectively. If they had responded within the deadline, they received a central green “*” (font size 20 px, 0.56°; 38 cd/m^2^) and otherwise a red “!” (font size 20 px, 0.56°; 22 cd/m^2^) as feedback (for 750 ms).

The participants’ task was to indicate whether a target stimulus, a black arrow (Figure 1, font size 20 px, 0.56°, < 1 cd/m^2^) pointed to the left or right. Participants of Experiment 1 performed a spatial Stroop task: The target appeared left or right (6°) to screen center. In the congruent condition, the target appeared on the side of its pointing direction, in the incongruent condition, it appeared on the other side. Participants of Experiment 2 performed a flanker task: The target appeared at screen center and was flanked left and right by two distractor arrows each. In the congruent condition, the distractors pointed in the same direction as the target, in the incongruent condition, in the other direction. In both experiments, participants were instructed to focus only on the target’s pointing direction and ignore its spatial location or the distractors, respectively. In addition, they were instructed to prioritize responding within the deadline, even if it meant to guess their response.

Every participant performed 1188 experimental trials in 5 sessions, yielding 5940 trials in total. All combinations of the fixation durations, gap durations, pointing directions of the target and congruency conditions occurred equally often and in randomized order within 9 blocks per session.

### Data analysis and eye movements

Data and analysis code are available on the Open Science Framework (https://osf.io/5rk6n/?view_only=4cc8ed616d434a8d91fe02b57a6d0908; private link for peer-review that will be made public upon manuscript acceptance for publication). Data were analyzed using R (4.1.0, https://www.R-project.org/). For each trial, the raw processing time (RPTs) was computed as reaction time (RT) minus gap duration (Salinas et al., 2019). Tachometric functions were derived by sliding a 1 ms bin across RPTs from −200 ms to 1000 ms and computing the average performance within the bin (Figure 2a, d). Trials with RPTs outside this range and were excluded from analysis (Experiment 1: 9.56%; Experiment 2: 21.19%). Performance (the proportion of correct trials) was then locally regressed on the RPT bins (using R’s loess-function with a span-parameter of 0.2) to obtain separate tachometric functions for the congruent and incongruent condition. This was done to analyze the congruent and incongruent conditions with the same function, even though they had qualitatively different shapes (*Figure 2a, d;* this step was different from Salinas et al.’s approach that focused specifically on the shape of the incongruent condition).

As the data consistent across participants, data analysis was performed on the data pooled across participants (following Salinas et al.’s analysis of their aggregate participant).

Experimental conditions were compared using permutation tests as follows. Original effects (e.g., differences between the congruent and incongruent condition in the minima of their estimated tachometric functions, see Figure 2b, d) were located in a distribution of effects from a 2000-fold reanalysis of the raw data with randomized condition labels, that is, trials labeled as congruent or incongruent.

## Acknowledgments

The author thanks Anika Krause for help with the data collection of Experiment 1 and 2, Niklas Dietze and Lukas Recker for comments on an earlier draft of this manuscript, and the team of the Neuro-cognitive Psychology Group at Bielefeld University for helpful discussions.

## Supplemental materials

### Pilot experiment

#### Results

Similar to Experiment 1, the pilot experiment assessed cognitive control using a variant of the spatial Stroop task (e.g., Lu and Proctor, 1995; Schneider, 2020). The experiment was similar to Experiment 1, but differed from it in a number of method details (see the methods below). The results of the pilot experiment are visualized in Figure S1. Tachometric functions were obtained and analyzed as for Experiment 1 (Figure S1a). As in Experiment 1, performance rose monotonously toward ceiling performance with increasing RPT in the congruent condition. In contrast, performance dropped below chance within a constrained time-window in the incongruent condition. This drop in the incongruent condition was statistically reliable, as indicated by a significantly lower minimum of the tachometric curve in the incongruent compared with the congruent condition (permutation test, *p* < .001, see Figure S1b).

**Figure S1.**
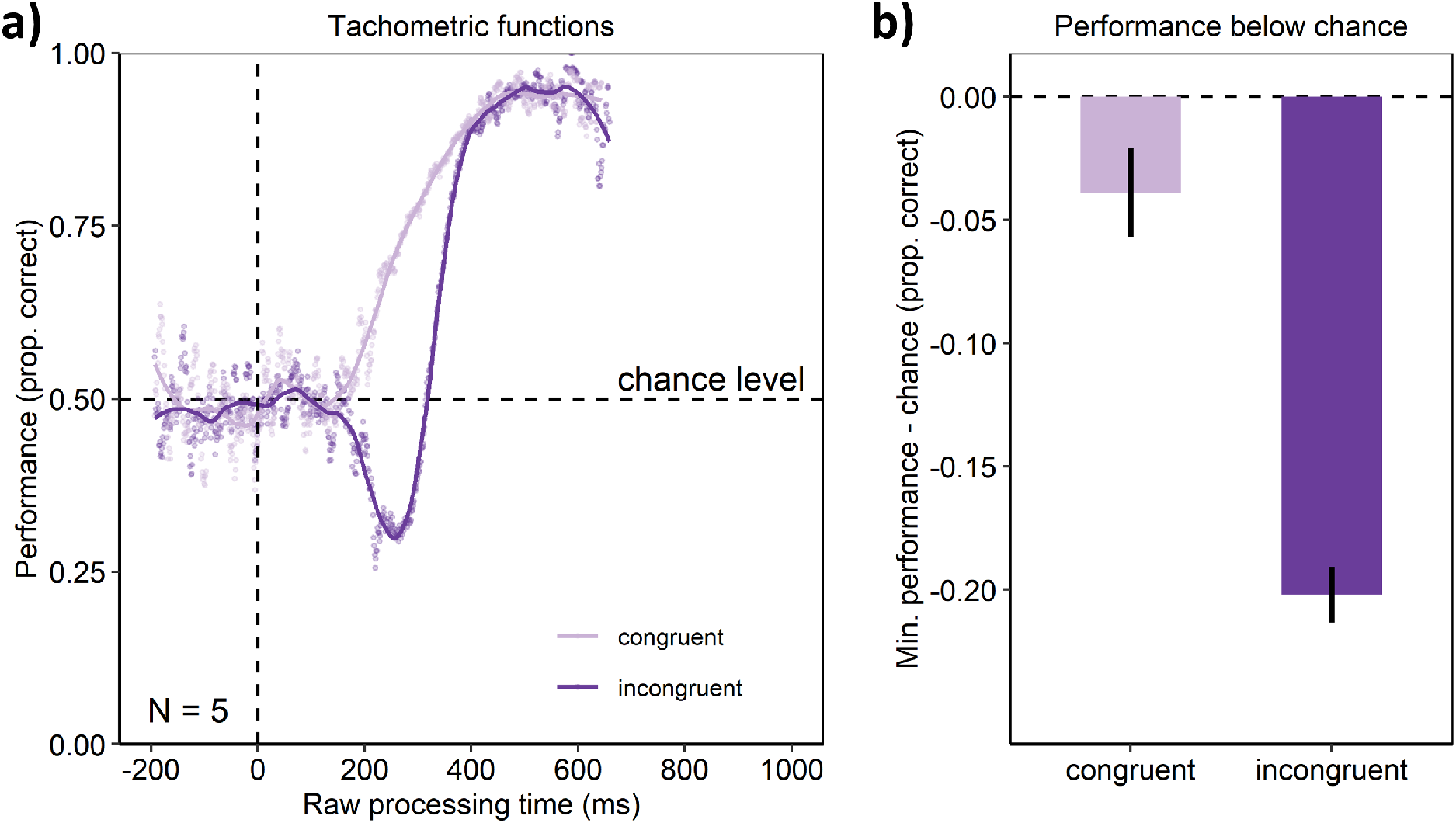
a) Behavioral results of the pilot experiment. a) Tachometric functions depicting performance as a function of RPT for the congruent and incongruent conditions of the aggregate participant. b) In contrast to the congruent condition, the incongruent condition led to a drop of performance below chance. Error-bars are standard errors from a bootstrap.

#### Materials and methods

##### Participants

Five participants (aged 20 – 29 years) took part in the pilot experiment (one additional participant did not perform the experiment, because the experiment did not run on their home PC, on which the experiment was performed, see the setup below). They all reported (corrected-to-) normal vision and gave written informed consent before participating. The experiment followed the ethical regulations of the German Psychological Society (DGPs) and was approved by Bielefeld University’s ethics committee.

##### Experimental setups and task

Participants performed the experiment at their homes using their own PCs (because laboratories were closed due to the COVID-19 crisis). They were instructed to set their screens at a refresh rate of 60 Hz (one participant’s screen had a variable refresh rate for technical reasons, with about 75 Hz on average) and to view them from a distance of 71 cm. Responses were collected using external computer mouses. The experiment was programmed using PsychoPy2 (v1.85.3; Peirce, 2008, 2007).

The task and the trial sequence was identical to the one of Experiment 1, with a few exceptions. Stimuli were shown against a black background. Target stimuli consisted of the letters “L” or “R” (Arial font, 1.5° height) on a white square (3.5×3.5°), appearing to the left or right (8° from screen center). The fixation stimulus was a small white square (0.1 × 0.1°). Participants performed 10 sessions of 568 trials each, in which all combinations of target letter (L vs. R), target location (left vs. right), fixation durations (350, 400, 500 ms), and gap durations (0, 100, 200, 300, 400, 500, 600, 700, 800, 900, 950 ms) occurred equally often.

##### Data analysis

The data analysis was identical to the corresponding analyses of Experiment 1.

### Physiological arousal in Experiment 1 and 2

The average pupil size within 1 s after the manual response (z-scored for the participant’s session) and the velocity of the first saccade (relative to its amplitude, detected using the Eyelink 1000 algorithm using a velocity threshold of 35°/s and an acceleration threshold of 9500°/s^2^) after target onset were analyzed as measures of physiological arousal (Di Stasi et al., 2013; Mathôt, 2018).

As can be seen in Figure S2, for both experiments, and for both, the congruent and incongruent condition, the two arousal measures monotonously increased with increasing gap duration. This indicates that introducing time-pressure for responding by this gap duration effectively changed participants’ momentary state of arousal. This should affect the overall responsiveness of the perceptual and cognitive brain systems, for example by modulating overall neuronal activity (e.g., by modulating the nor-epinephrine system, Aston-Jones and Cohen, 2005). Thus, taken together, the results from these physiological measures indicate that the manipulation successfully introduced an internal state of urgency.

**Figure S2.**
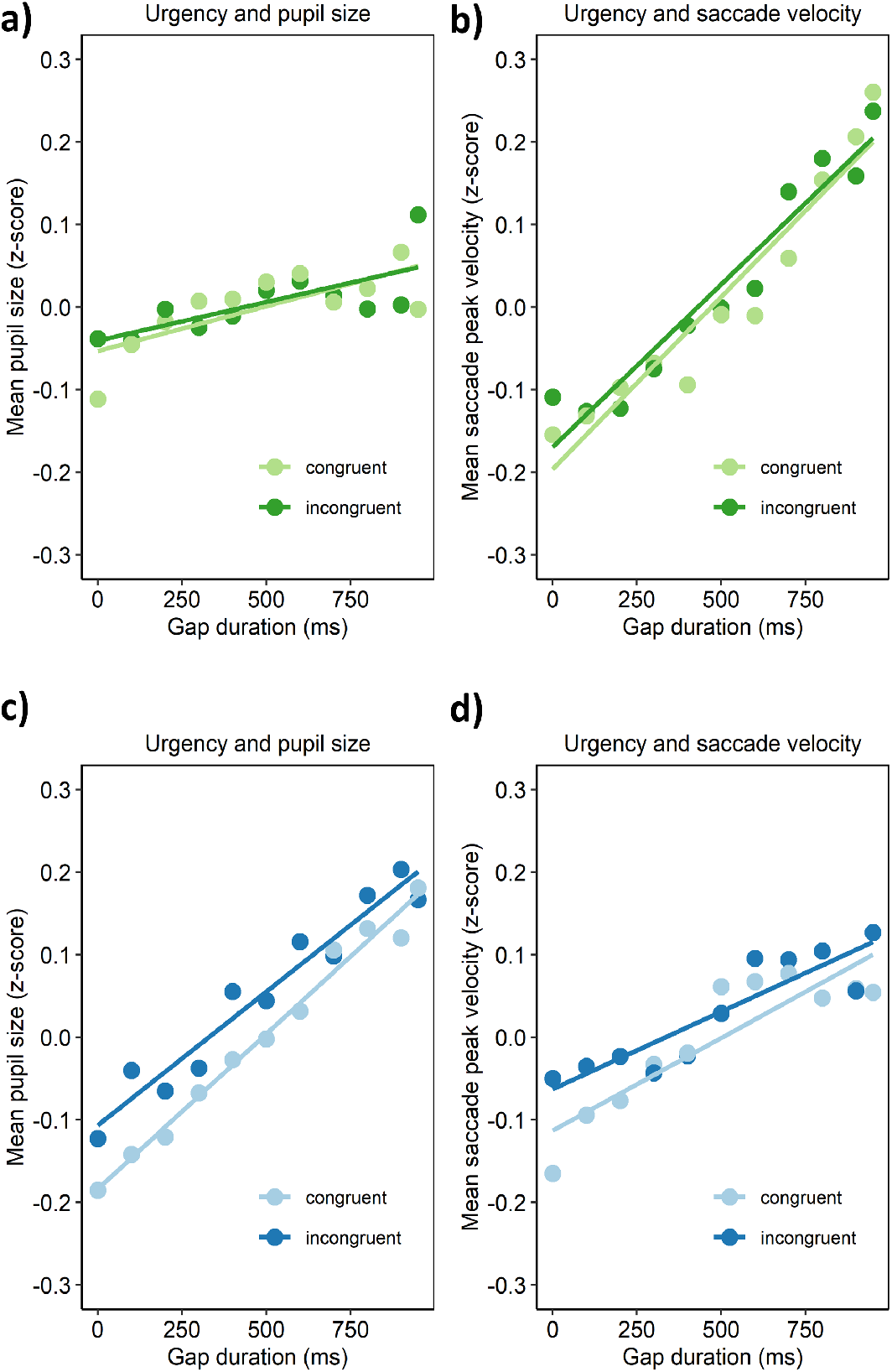
Relationships between urgency and measures of arousal in Experiment 1 (upper panel) and Experiment 2 (lower panel). a/c) Urgency (gap duration) was associated with pupil size both congruency conditions (average pupil size was assessed over 1 s after manual responses and z-scored across the trials of each session, see methods). b/d) Likewise, urgency was associated with the peak velocity (relative to saccade amplitude) of the first saccade after target onset in both congruency conditions (see methods for saccade detection criteria).

